# Gene-gene relationships in an *Escherichia coli* accessory genome are linked to function and mobility

**DOI:** 10.1101/2021.03.26.437181

**Authors:** Rebecca J. Hall, Fiona J. Whelan, Elizabeth A. Cummins, Christopher Connor, Alan McNally, James O. McInerney

## Abstract

The pangenome contains all genes encoded by a species, with the core genome present in all strains and the accessory genome in only a subset. Coincident gene relationships are expected within the accessory genome, where the presence or absence of one gene is influenced by the presence or absence of another. Here, we analysed the accessory genome of an *Escherichia coli* pangenome consisting of 400 genomes from 20 sequence types to identify genes that display significant co-occurrence or avoidance patterns with one another. We present a complex network of genes that are either found together or that avoid one another more often than would be expected by chance, and show that these relationships vary by lineage. We demonstrate that genes co-occur by function, and that several highly connected gene relationships are linked to mobile genetic elements. We find that genes are more likely to co-occur with, rather than avoid, another gene, suggesting that cooperation is more common than conflict in the accessory genome. This work furthers our understanding of the dynamic nature of prokaryote pangenomes and implicates both function and mobility as drivers of gene relationships.

**Data summary:** All Supplementary Data files and the Python scripts used in the analyses are available at doi.org/10.17639/nott.7103.

**Impact statement:** The pangenome of a species encompasses the core genes encoded by all genomes, as well as the accessory genes found in only a subset. Much remains to be understood about the relationships and interactions between accessory genes; in particular, what drives pairs of genes to appear together in the same genome, or what prevents them from being in the same genome together, more often than expected by chance. How these co-occurrence and avoidance relationships develop, and what effect they have on the dynamics and evolution of the pangenome as a whole, is largely unknown. Here, we present a springboard for understanding prokaryote pangenome evolution by uncovering significant gene relationships in a model *Escherichia coli* pangenome. We identify mobile genetic elements and the sharing of common function as possible driving forces behind the co-occurrence of accessory genes. Furthermore, this work offers an extensive dataset from which gene relationships could be identified for any gene of interest in this *E. coli* accessory genome, providing a rich resource for the community.

## Introduction

*Escherichia coli* is one of the most widely used and studied bacterial species in microbiology. Recent efforts have resulted in a cataloguing of the essential genes in this species using transposon-directed insertion site sequencing (TraDIS) (1), thereby defining which genes are indispensable and which are not. It is arguable that the accessory genes found in the *E. coli* pangenome are not essential for the survival of the species, making it somewhat of a curiosity that such a large set of accessory genes is maintained. A separate study of a dataset of 53 *E. coli* genomes identified more than 3,000 metabolic innovations that all arose as a consequence of the acquisition via horizontal gene transfer (HGT) of a single piece of DNA less than 30 kb in length, and that 10.6% of innovations were facilitated by earlier acquisitions (2). This suggests therefore that HGT has the ability to bring together sets of genes that, when combined, provide benefits over and above the benefits that the genes could confer on their own. Indeed, it is likely that there are situations where the genes on their own might be deleterious, but in combination they confer a fitness advantage (3), thereby contributing to the maintenance of a large accessory genome.

Analyses of genome content shows that *E. coli* pangenome evolution is driven by differential gain and loss of accessory genes, including plasmids, phage, and pathogenicity islands (4; 5; 6). What is not yet clear is the underlying structure of the pangenome and which genes are key influencers of the presence or absence of other genes in a given genome. Plasmid mobility, for instance, is anticipated to be an important agent of pangenome structuring (7). Plasmids engage a diverse range of proteins for the purpose of plasmid partitioning and maintenance, but the presence and absence of these protein encoding genes in accessory genomes has not been established concretely.

Variation in overall genome sequence has the potential to influence which other genes can or cannot be present in a genome. A gene might be beneficial to one strain of a species, but deleterious in a different strain. Sets of genes encoding proteins that together form an essential biosynthetic pathway, for example, are expected to co-occur in the same genome, likewise those that form multi-protein complexes. The presence of a gene or set of genes can also exclude, or ‘avoid’, others from the same genome. Competitive exclusion is exemplified in *Salinispora* species, where only one of two iron-chelating siderophore gene clusters is ever found in a given strain, despite evidence of frequent HGT (8). An understanding of gene-gene dependencies and influences can therefore provide a real insight into how pangenomes originate and are maintained. We can address at least two questions when we look at gene-gene co-occurrence and avoidance patterns. We can ask whether the pangenome has been structured by random genetic drift or by natural selection, and we can ask whether gene co-occurrence has been more important than avoidance in a pangenome.

In practical terms, the large *E. coli* accessory genome can serve as a testbed for calibrating how much gene co-occurrence and avoidance can teach us about pangenome origins and evolution. A considerable amount of diversity is found across all known sequence types (STs) (4; 9; 10; 11; 12), while a large body of experimental validation of gene function has had the effect of reducing the number of unknown open reading frames (ORFs) in the pangenome. It is anticipated that there may be collections of lineage-specific coincident gene-gene relationships, but the process driving these relationships is not yet clear.

Here, we interrogate *E. coli* pangenome dynamics in order to identify coincident gene-gene relationships (13). In a large sample of the total *E. coli* pangenome, comprising 400 judiciously selected genomes spanning 20 different STs, we have identified significant gene-gene co-occurrences and avoidances. We find connected components that are enriched in genes that share a common function, suggesting that the role that genes play in a system can partially explain their significant co-occurrence. We also find that MGEs extensively influence gene co-occurrence and avoidance, including for genes linked to detoxification and antimicrobial resistance (AMR). We identified more co-occurrence relationships than avoidance relationships, which, according to our data and method of analysis, suggests that gene cooperation is more common than conflict in the pangenome. Our results provide an extensive set of possible empirical experiments that, together with our *in silico* predictions, further our understanding of the complex ecological interactions and dynamics in the *E. coli* pangenome.

## Methods

### *E. coli* genomes and pangenome

A set of 400 *E. coli* genomes were downloaded from EnteroBase using a custom Python script (github.com/C-Connor/EnterobaseGenomeAssemblyDownload). The set contained 20 genomes from 20 different STs; ST3, ST10, ST11, ST12, ST14, ST17, ST21, ST28, ST38, ST69, ST73, ST95, ST117, ST127, ST131, ST141, ST144, ST167, ST372, ST648. The STs were chosen to include multiple extraintestinal pathogenic *E coli* (ExPEC), enterohemorrhagic *E. coli* (EHEC), and commensal lineages. The sequences were annotated using Prokka (14) and a gene presence-absence matrix was generated with Panaroo (15) using the default settings and the -a core flag to generate a core gene alignment.

### Core gene phylogeny

The core gene alignment file produced by Panaroo was trimmed using trimAl (16) with a -gt value of 1. Phylogenetic relationships between the strains were inferred using the core gene set from the pangenome. A core gene phylogeny was constructed from the trimmed alignment using the IQ-Tree software (17) with the GTR+I+G substitution model. Phylogenetic tree visualisation was carried out using the Interactive Tree of Life v5 (iTOL) (18).

### Coincident gene identification and visualisation

Gene co-occurrences (associations) and avoidance (dissociations) were identified using Coinfinder (13) using a threshold of 0.01 for low abundance filtering and employing the Bonferroni correction to account for multiple testing. Networks were visualised using the Gephi software program (v0.9.2) with the Fructerman-Reingold algorithm used for graph layout. Heatmaps were generated using seaborn (v0.10.1). Hub genes are identified as genes forming a number of gene co-occurrences or avoidances that is 1.5 times the interquartile range (IQR), as in (19). Gene names were taken from the Panaroo gene cluster identifications.

### Prediction of plasmid-encoded genes

Genes were determined to be chromosomal or plasmid-encoded by collecting each contig containing the gene across all strains in which it was present, and then assessing whether those contigs were predicted to be plasmid- or chromosome-derived sequences using mlplasmids (20).

### Phosphotransferase system analysis

A list of all phosphotransferase system (PTS) genes was obtained from the KEGG database (21) and the *E. coli* gene presence-absence matrix was used to highlight those genes found in the *E. coli* pangenome.

### Data availability

All Python scripts used for data analysis and figure construction are freely available at doi.org/10.17639/nott.7103. The gene presence-absence matrix generated by Panaroo, core gene phylogeny, outputs from Coinfinder, and a list of genomes mapped to their ST are also available as Supplementary Data at the same location. Descriptions of Coinfinder outputs can be found in the original software publication (13).

## Results

### The *E. coli* pangenome is highly structured

An *E. coli* pangenome, composed of 400 genomes from 20 different STs, was constructed (Fig. S1). Panaroo was used for this analysis as preliminary investigations found that the clustering produced fewer incidences of false positives in the avoidance network than when the gene presence-absence matrix was constructed using Roary (22). This pangenome sample consists of 3,191 core, 120 soft core, 2,935 shell, and 11,665 cloud genes (Fig. S2), replicating previous observations that *E. coli* has an open pangenome (9). Using Coinfinder (13) we identified a gene co-occurrence network that consisted of 8,054 nodes joined by a total of 500,654 edges (Fig. 1a), and an avoidance network consisting of 3,203 nodes joined by 203,503 edges (Fig. 1b). The co-occurrence network is therefore larger, both in terms of numbers of nodes and also numbers of interactions, though both networks have similar average numbers of connections per node. Of all gene clusters in the gene presence-absence matrix, 45.0% form at least one co-occurrence pair, and 17.9% at least one avoidance. Of the accessory genes analysed by Coinfinder, 77.8% form at least one co-occurring or avoidant pair.

**Fig. 1.**
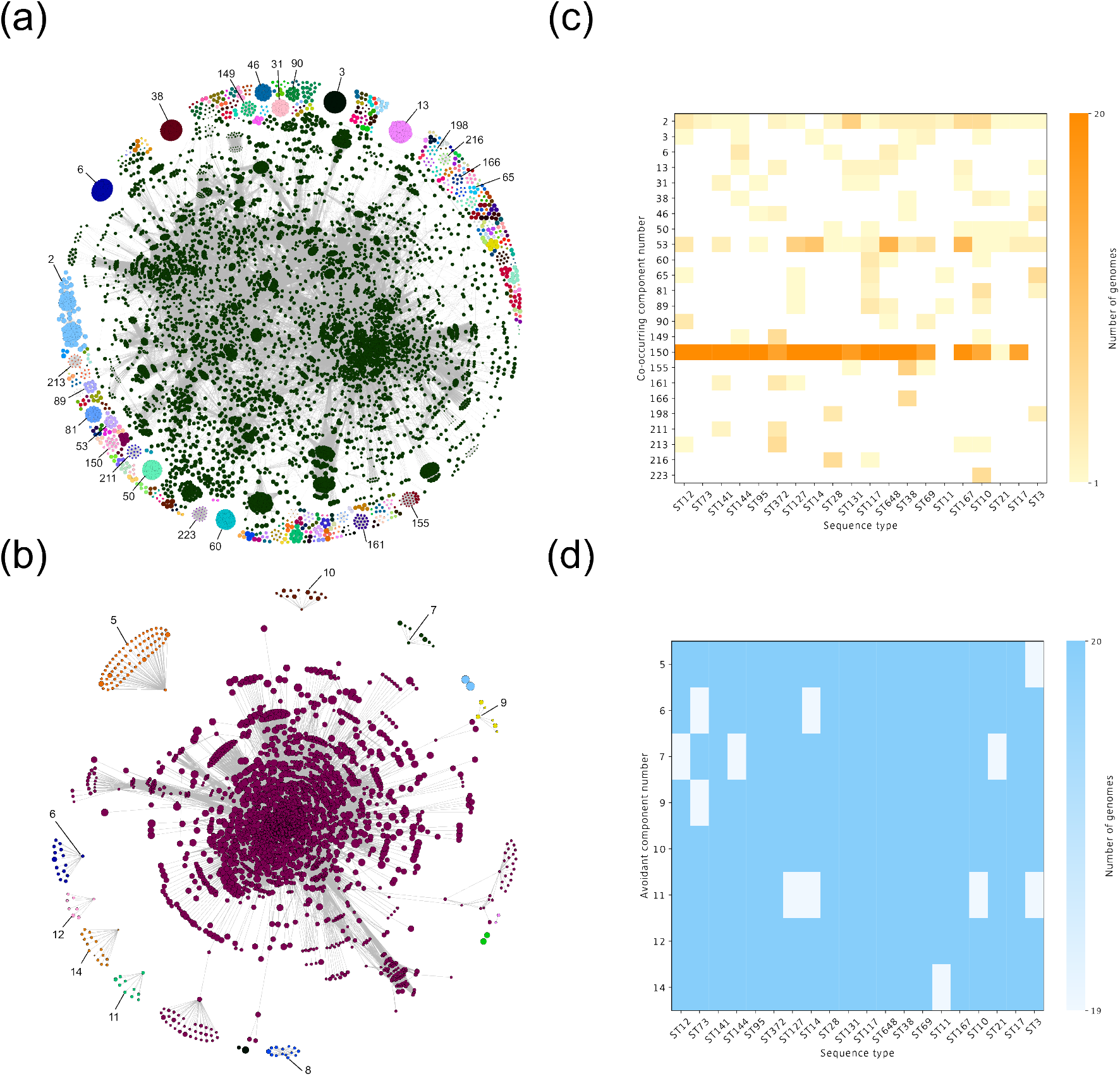
Gene relationships in the *E. coli* pangenome. (a) An overview of the co-occurrence and (b) avoidance networks, coloured by connected component. Selected connected components are numbered as in the Supplementary Data Files. (c) The number of genomes in which part or all of a co-occurrence and (d) avoidance connected component appears in each ST. An absence of a co-occurrence component in a ST is depicted as colourless.

### Co-occurring genes share function

The co-occurrence network contains 224 connected components, including one large connected component with extensive interactions (Fig. 1a). If indeed co-occurrence is shaped by functional interactions and dependencies, we could expect co-occurrence analyses to pick out known subsystems. Component 150, which is made up of a total of 15 genes, consists in part of twelve genes involved in the type II secretion system (T2SS) (*epsC-H, epsLM, gspK, pppA* and *xcpVW*) that facilitates the translocation of a wide variety of proteins from the periplasm to the exterior of the cell. Similarly, component 50 includes *epsE* (T2SS), and the type IV secretion system (T4SS) genes *vir1,4,8,10,11*. Component 150 has a broader distribution across the STs (n=18) than component 50 (n=6), and is found at a much higher abundance. In fact, for 11 of the STs, all 20 genomes contain component 150 (Fig. 1c). These data illustrate that indeed analyses of co-occurrence patterns, as implemented by Coinfinder, can pick out functional dependencies.

The constituent genes in components 3, 13, 60, and 149 are outlined in Table 1 and are known to function in DNA replication. Of note in these components are the plasmid copy control gene *repB* and the plasmid partitioning protein-encoding gene *parB*, both known to be plasmid-encoded, as well as the chromosome-encoded *dnaJ* that functions in plasmid replication (23; 24). The fact that these functions are found across four separate co-occurrence components shows that within this set of functions related to mobility, there are co-occurring communities consisting of distinct groups of genes that preferentially co-occur. These components are found in a variety of STs, though in low abundance (Fig. 1c).

**Table 1.** Certain co-occurrence components function in fundamental cellular processes. The known gene clusters relating to DNA and RNA replication, regulation and repair found in the co-occurrence connected components 3, 13, 60, and 149. Genes are named as per the Panaroo gene clusters in the gene presence-absence matrix. Where the gene is identified by ‘group_’, a gene identifier, taken from the Panaroo non-unique gene name, is given in parentheses.

| Component 3 | Component 13 | Component 60 | Component 149 |
| --- | --- | --- | --- |
| dnaB_2_dnaB_1_dnaB_3 | polA_2 | dnaB_3_dnaB_2 | group_4304 (*recQ*) |
| dnaJ_3_dnaJ_1_dnaJ_2 | dnaE_1_dnaE_2_dnaE1 | ssb_1_ssb_5_ssb_4 | srmB_2 |
| ssb_3_ssb_2 | dnaG_1 | group_3992 (*topB*) | group_296 (*rapA*) |
| smc__smc_1 | dnaQ_1_dnaQ_2 |  |  |
|  | group_7368 (*parB*) |  |  |
|  | repB_2 |  |  |
|  | lig |  |  |
|  | smc_2_smc |  |  |

### Co-occurrence gene hubs are linked to virulence and mobile elements

We found considerable variation in the number of significant co-occurrence or avoidance relationships for any individual gene in the *E. coli* accessory genome. From the degree distribution in the co-occurrence network we identified a large group of 427 highly connected “hub genes” (Fig. 2a), with a high of 896 co-occurrences recorded for an individual gene cluster (group_2839, identified as dnaC_4 from the non-unique gene name). These hub genes either facilitate or promote the existence of a large number of other genes in any given genome. The most common known functions of co-occurrence gene hubs are attack and defence (including toxin-antitoxin systems, Shiga toxin, CRISPR system subunit, and genes encoding hemolysin and proteins involved in detoxification), metabolism, and DNA and RNA processes (including DNA primase, helicase, and replication proteins, tyrosine recombinases, and proteins involved in DNA repair) (Fig. 2b). Full lists of the number of pairs formed by individual genes are provided as Supplementary information.

**Fig. 2.**
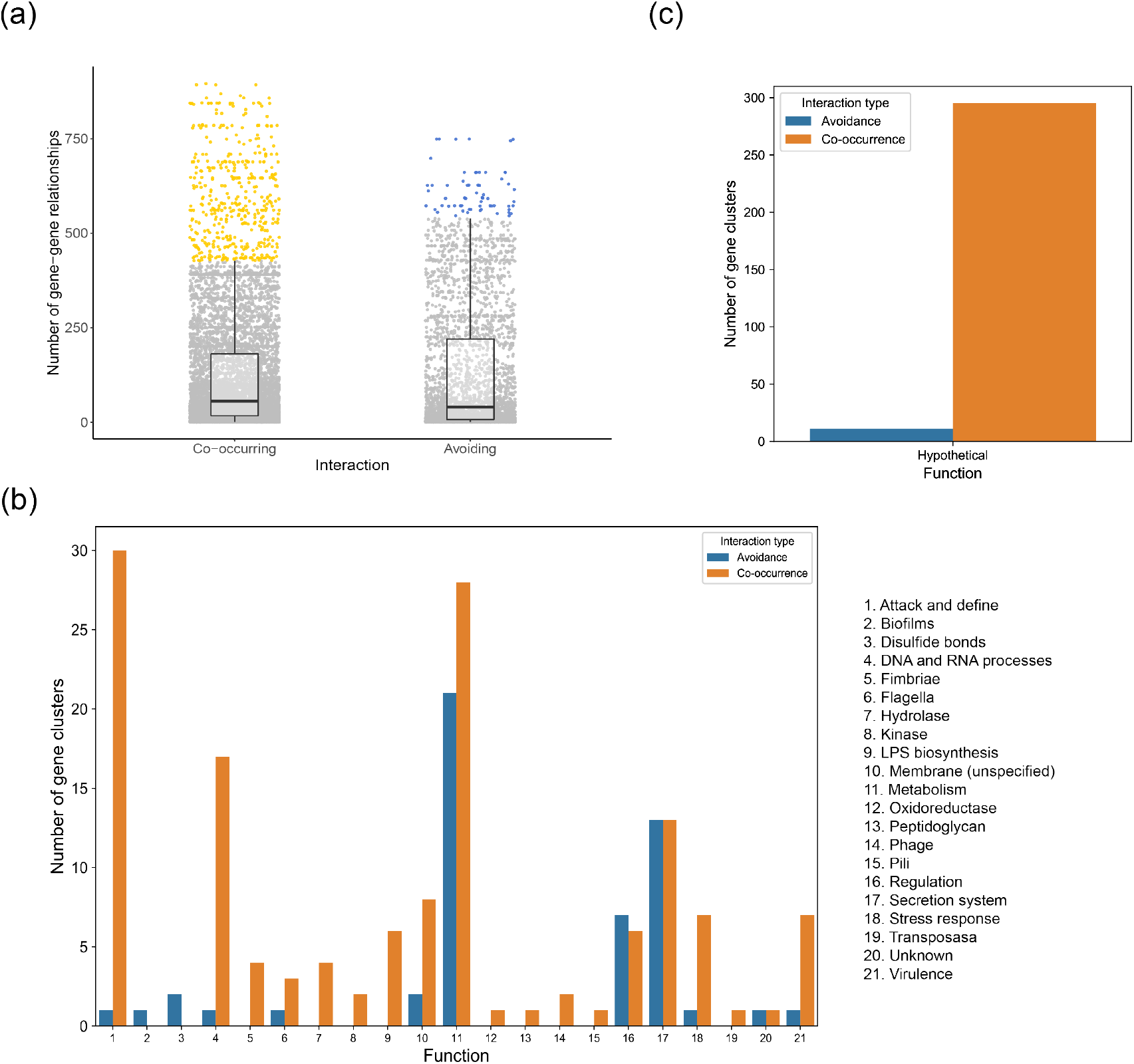
Highly connected hub genes in the accessory genome. (a) The number of gene-gene relationships formed by individual genes. Hub genes (1.5 times the upper IQR) are coloured orange (co-occurrence) and blue (avoidance). (b) Co-occurrence (orange) and avoidance (blue) hub genes categorised by loose biological function. Attack and defence includes CRISPR system subunits, toxin-antitoxin systems, toxin production, detoxification. DNA and RNA processes includes replication, repair, recombination, and plasmid segregation. (c) The number of hub genes encoding hypothetical proteins.

A notable subset of the hub genes is linked to DNA exchange and MGEs. Some are known plasmid-encoded genes (*toxB, hlyACD*, and *ssb* (24; 25)), and the Shiga toxin subunits *stxAB* (7) are encoded on a single phage (26). The tyrosine recombinases *xerCD* function in plasmid segregation, as does *parM*, and there are several genes either prophage-encoded (*recE* (27)) or related to phage functions (*intQ, tfaE*). The transposase *tnpA* is also a hub gene, forming a co-occurring pair with 841 other genes. These findings indicate a process where hub genes that have a role in lateral mobility of genetic material are specifically co-occurring with genes that confer fitness advantages, on average, when mobilised. This suggests a process of mutual benefit; the mobility enablers co-occurring with the genes most likely to confer positive fitness effects.

### The avoidance network is characterised by root excluders

The avoidance network has substantially fewer connected components (n=14) than the co-occurrence network, and many of these components are characterised by the presence of a single “root excluder”; a situation where one gene avoids a gene set that in turn all co-occur with one another (Fig. 1b, Fig. S3). Over half of the components in this network show this pattern, specifically components 5-7, 9-12, and 14 (Table S1). These components are found in at least 19 genomes of all STs (Fig. 1d). None of the root excluders form avoidance hubs (Fig. 2a). As an example, within component 14, the root excluder *dhaR*, a transcriptional regulator of the dihydroxyacetone kinase operon (28) avoids 11 genes involved in the production of T2SS proteins (*gspHK, epsEFLM, pulDG, outC, xcpVW*), amongst other known and hypothetical genes. The plasmid-associated nature of some T2SS genes (24) suggests an active process of dissociation or avoidance as a result of their mobility.

### Avoidance hub genes are lineage-specific

The avoidance network is smaller than the co-occurrence network with fewer hub genes, though high degree nodes can be identified with the highest showing 749 avoidances for a single gene cluster (atoA, atoD, zraR_2_spo0F, and zraS_2) (Fig. 2a). The *ato* genes encode proteins involved in the degradation and transport of short-chain fatty acids (29), and the *zra* genes encode a two-component system linked to antimicrobial tolerance in *E. coli* (30). The most enriched function of the avoidance hub genes is in metabolism, and when grouped by function, regulation is the sole category where the number of avoidance hubs is greater than the number of co-occurrence hubs (Fig. 2b).

Some genes avoid considerably more genes than they co-occur with. For instance, the putative diguanylate cyclase *ycdT*, which catalyses the production of cyclic di-3’,5’-guanylate and is reportedly under positive selection in uropathogenic *E. coli* (12), significantly co-occurs only with *pgaA-D*. The proteins encoded by *pgaA-D* are involved in biofilm formation. In contrast, *ycdT* avoids 83 other gene clusters including the CRISPR system Cascade subunits *casACDE*. All 400 genomes have at least one of either *ycdT* or the *casACDE* genes, but they are found in isolation in 298 genomes. The only STs where this *ycdT*-*casACDE* avoidance pattern is not observed in any of the genomes are ST3, ST17, ST11, ST38, and ST128, and therefore does not demonstrate a clear phylogenetic split (Fig. S1). We might speculate that there is another element of genome variation that modulates whether *ycdT*-*casACDE* can or cannot be present in the same genome.

The avoidance hub genes are also enriched in functions related to secretion systems (Fig. 2b). We found that between 627 and 661 gene clusters avoid the T2SS-related genes *gspA-M* and the probable bifunctional chitinase/lysozyme that is secreted by the T2SS (31). Of these, one is particularly pertinent; *spiA* (also known as *escC*), encoding a type III secretion system (T3SS) outer membrane secretin (32). Incidences of *spiA* avoiding *gspA-M* gene clusters are found in all 20 STs. These data provide some evidence that secretion systems may have a substantial genome-wide influence.

### Repeated co-occurrence of resistance genes and transposon genes

We have demonstrated that genes that share functions or pathways will commonly co-occur, and that certain highly connected co-occurrences and avoidances involve genes associated with plasmids. We hypothesised that other agents of horizontal transfer could therefore form the centre of co-occurrence hubs as a result of their mobility. The transposase *tnpA*, for example, is a co-occurrence hub gene (n=841 gene pairs) (Fig. 2a). This is the only transposon of the 22 within the co-occurrence network that is classified as a hub gene. We found that three connected components (excluding the large component in the centre of Fig. 1a) contain transposons.

The first, component 81, is comprised of a collection of genes enriched in functions related to metal detoxification. Eight genes relate to copper (*copABDR, pcoCE*) and silver (*silPE*) resistance, and six encode proteins involved in producing a cation efflux system (*cusABCFSR*) (Fig. 3a). A Tn7 transposition protein, *tnsA*, is also present. These systems may be acquired and retained together in order to provide a broader spectrum of metal detoxification. This connected component is only found in four of the 20 STs; ST3 (number of individual genomes = 2), ST10 (n=5), ST117 (n=2), and ST127 (n=1) (Fig. 1c). These STs are not close phylogenetically (Fig. S1), indicating that it is not simply a lineage dependent characteristic.

**Fig. 3.**
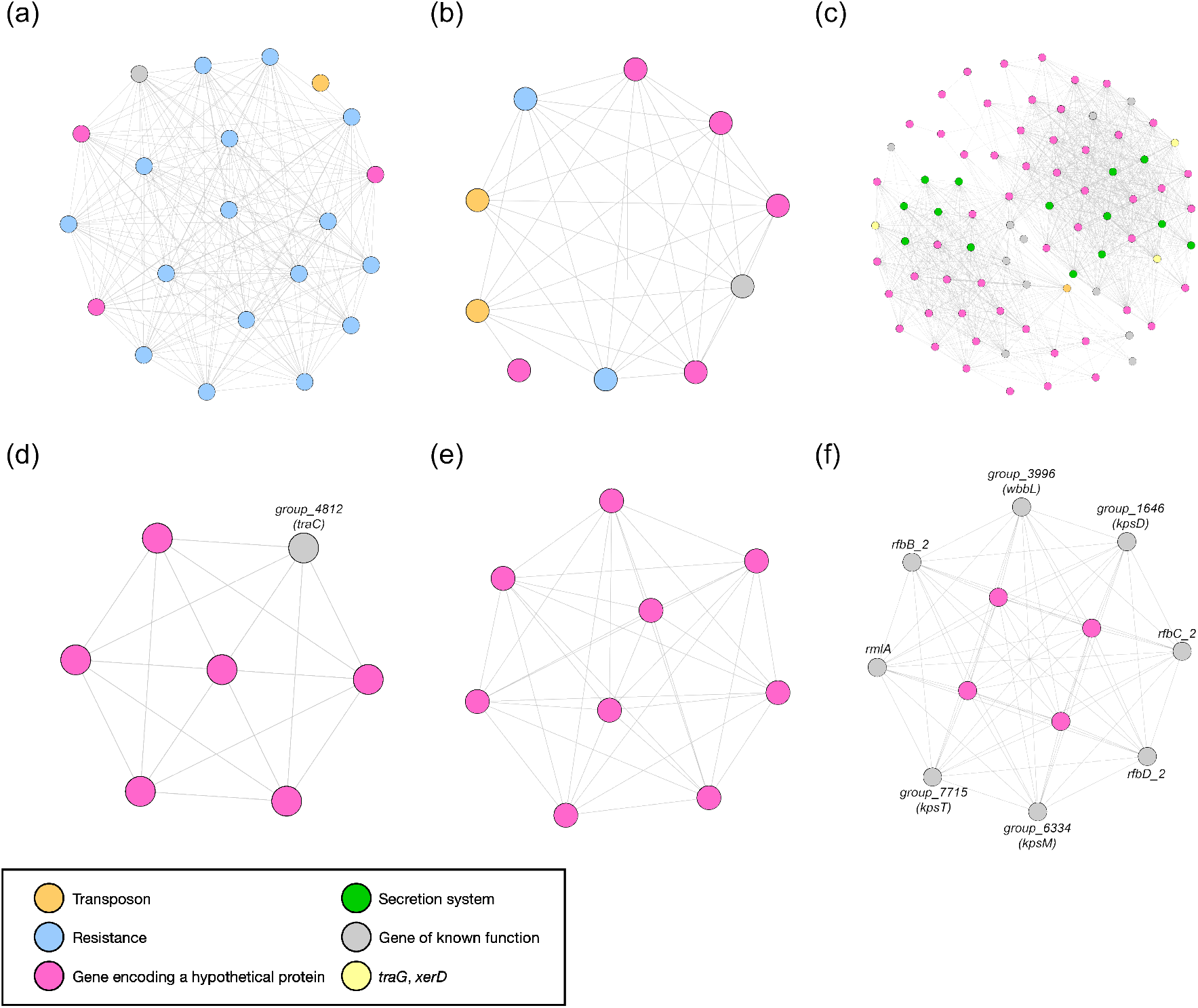
Select co-occurrence connected components are linked to transposons and ones enriched in genes encoding hypothetical proteins. (a) Co-occurrence component 81, enriched in genes related to metal detoxification. (b) Co-occurrence component 53, linked to tetracycline resistance. (c) Co-occurrence component 2, comprised in part of genes encoding secretion systems. (d) Co-occurrence components 198 and (e) 89 are enriched in genes encoding hypothetical proteins. (f) Co-occurrence component 90 contains genes relating to capsule formation. Select gene clusters are labelled.

Component 53 consists predominantly of the Tn10 transposon proteins *tetCD* that confer tetracycline resistance, as well as *gltS* (a sodium/glutamate symporter) and five hypothetical proteins (Fig. 3b). This component is found in more than half of the genomes in the multidrug resistance-associated ST648 (n=13) and ST167 (n=12), and in at least one genome in 16 of the 20 STs (Fig. 1c). The co-occurrence of *gltS* and the hypothetical proteins with the tetracycline resistance genes could suggest they may either be linked in a novel way to AMR, or, alternatively, that they have been transferred with the resistance genes and their co-occurrence is purely as a result of this hitchhiking effect.

The remaining connected component that contains a transposon is component 2. This component is comprised of two quasi-cliques and consists, in part, of the T4SS genes *virB1,2,4,6,8-11*, a putative transposon Tn552 DNA invertase (*bin3*), the conjugal transfer protein *traG*, and the tyrosine recombinase *xerD* (Fig. 3c). The presence of these genes throughout both of the quasi-cliques strongly suggests that MGEs are influencing the co-occurrence relationships in this component. This component is found in at least one genome in 18 different STs (Fig. 1c), providing further evidence for gene mobility. The transposons could influence the transfer of the genes in these components, or they could be hitchhiking between genomes alongside the rest of the component.

### Hypothetical proteins form a hidden network with a large influence on the *E. coli* accessory genome

A theme that emerged within the connected components is the central role in structuring the pangenome that is clearly being played by many genes with unknown function. First, 67.5% of the co-occurrence hub genes (n=295) and 17.5% (n=11) of the avoidant hub genes encode hypothetical proteins (Fig. 2c). This was surprising given the extent to which *E. coli* has been studied. While many genes in our dataset have assigned functions, 11,491 ORFs in the gene presence-absence matrix used as input to Coinfinder were not ascribed a function, 64.2% of the total. Second, certain connected components are enriched in genes encoding hypothetical proteins, with several connected components consisting solely of unknown ORFs. Examples of this are co-occurrence components 198 and 89 (Fig. 3d, e), both of which are found in several different STs, though no ST contains both components (Fig. 1c). The entirety of component 89 consists of hypothetical proteins, while all but one of component 198 is a hypothetical protein. It should also be noted that both of these connected components also form a clique; every node in the connected component is connected to every other node. This means that every gene in the component shows a significant co-occurrence pattern with every other gene. It is therefore highly likely that these groups of genes are tightly, functionally linked to one another, though there is no published information related to the function of any of these genes.

We observe several connected components where the majority of genes in a component share a similar function. This makes it tempting to imply a putative role for the genes encoding the remaining hypothetical proteins in those components. Co-occurrence component 90 consists of five genes with known functions; three genes that are found in the dTDP-rhamnose biosynthesis pathway, a rhamnoslytransferase, and a polysialic acid transporter. This leaves four genes in this connected component that encode hypothetical proteins (Fig. 3f). We could speculate that these hypothetical proteins may therefore function in lipopolysaccharide or capsule formation (33). Component 2 consists of 82 gene clusters, including 53 that encode hypothetical proteins and 14 that relate to secretion systems (Fig. 3c). These observations show that there is an important network of genes of unknown function that exert a large influence on the *E. coli* accessory genome, and that our approach can generate meaningful hypotheses to inform investigation of the function of those genes.

### The drivers of gene-gene relationships are multifaceted

We have identified co-occurrence connected components that entirely consist of, or are enriched in, genes that share a common, identifiable role. We have also highlighted gene relationships that appear to be influenced by the presence of MGEs. If function and mobility were the only drivers behind significant gene-gene relationships, it might be expected that genes encoded on the same operon, or that function in a discrete system, will share the sames patterns of co-occurrence. To test this, we focused on phosphotransferase system (PTS) genes, which are used to import carbohydrates (34). These systems were chosen because several have been identified in *E. coli*, they are typically multi-component (35), and the presence of one system might logically be linked to the presence or absence of another.

We collated all genes encoding PTS components and identified those that form a co-occurring pair with at least one other PTS gene. We found no correlation between whether the encoded proteins are membrane-bound or cytoplasmic and the likelihood that they will co-occur with another PTS gene. Of the 67 PTS genes detailed in the KEGG database (21), 29 were not observed in our gene presence-absence matrix, and a further ten did not manifest a significant co-occurrence with any other PTS gene (Fig. 4a).

**Fig. 4.**
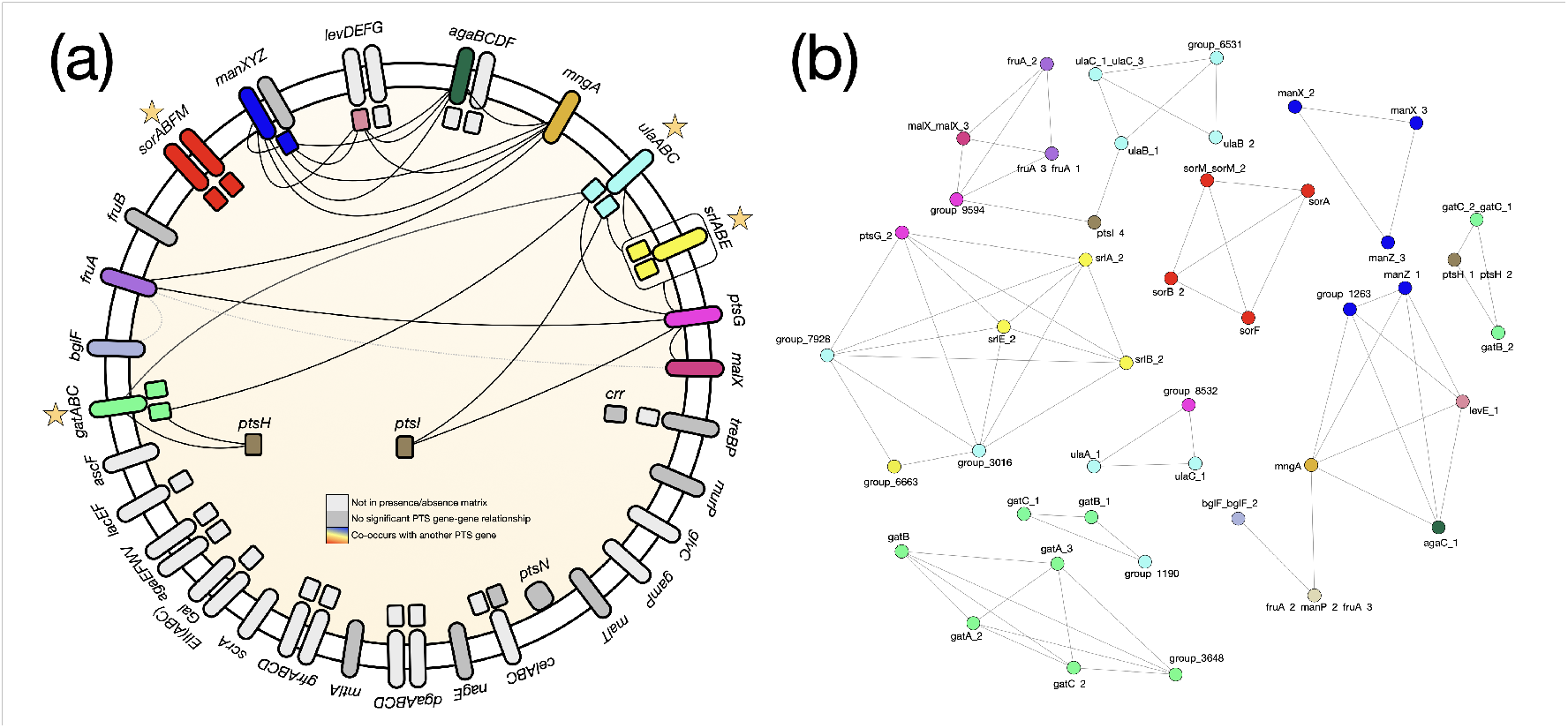
PTS gene relationships are not universal. (a) A schematic overview of the co-occurrence PTS gene relationships. PTS gene systems found in the KEGG database are coloured by those not present in this *E. coli* pangenome (white), those which do not form a coincident PTS-PTS pair in the accessory genome (grey), and the clusters that form a co-occurring pair with another PTS gene (coloured by system). A significant relationship is indicated by a solid black line. Where a relationship is observed with all genes in a PTS, the line connects to a box around the system. Transporters that consist of genes that all significantly co-occur with one another are marked with a yellow star. The grey dashed line connecting *fruA* with *bglF* and *malX* indicates a significant relationship in both the co-occurrence and avoidance datasets for different clusters of the same gene identification. (b) The specific co-occurrences by gene cluster. Nodes are coloured by system as in (a).

We found a complex pattern of co-occurrence across the systems. Certain PTS genes do show a consistent pattern for the complete system. For example, *srlABE* all co-occur with the same genes, and *sorABFM* only co-occur with each other (Fig. 4a, b). In contrast, the *manXYZ, levDEFG, agaBCDF*, and *ulaABC* gene relationships are not system-specific, varying instead by individual gene. Selection pressures on these systems are complex and heterogeneous, and gene co-occurrence relationships are likely driven by multiple factors.

## Discussion

The open pangenome of *E. coli* (9) provides a rich testbed to help understand genome and pangenome dynamics. Factors such as the immediate microenvironment in which a strain lives are known to influence the accessory gene content of any given genome and, consequently, the pangenome (36; 37; 38). Though there are known examples of how the fitness of one gene in a genome is influenced by the presence or absence of other genes, there has been no systematic, large-scale study of how genetic background, in terms of the presence or absence of genes in a genome, influences the fitness effect of an incoming gene (13). Recently, however, it has been shown that the genetic background of a genome has a direct effect on whether or not a gene is essential (39). We have little knowledge of why some lineages encode genes that others do not, and the extent to which the observed encoded genes influence the likelihood of successfully integrating other incoming genes. To begin to unpack this, we have presented a network of coincident gene relationships in a model *E. coli* pangenome. We found that nearly half of all gene clusters in the pangenome form a significant pair, and that this relationship is more likely to be one of co-occurrence than avoidance. This indicates an ecological relationship between genes in that cooperation is more likely than conflict; genes co-occur because they share function, whereas direct, antagonistic avoidances are less common. This shows that the *E. coli* pangenome is highly structured.

Previously, little was known about structure in prokaryote pangenomes; whether pangenomes arise as a result of drift, or whether they are maintained by selection (40; 41; 42). Recent genus-level analysis of a *Pseudomonas* pangenome found significant co-occurrence of genes that share common function, providing support for selection as a driver of pangenome evolution (19). We build upon this work with the observation that many co-occurring connected components are enriched in genes that share function, broadly defined. We also propose, alongside the role that sharing function plays, that gene mobility could be an additional mechanism behind the formation of these gene-gene relationships. MGEs are a known link to gene essentiality and virulence in *E. coli* (4; 39; 43), and they have been implicated in driving accessory genome differences in a *Listeria monocytogenes* pangenome (44). Here, we have demonstrated that they also influence gene co-occurrences by uncovering hub genes and connected components linked to or encoded on MGEs (24; 25; 7; 45; 46; 47; 48; 49). This includes known phage-encoded virulence factors (26; 50), transposons, and genes involved in conjugation. Together, this progresses current understanding of prokaryote pangenomes by suggesting that they are structured and dynamic, but also further underscores the importance of HGT in driving pangenome evolution (51).

Furthermore, we found that these gene relationships are often non-randomly distributed across the pangenome, with some being more frequently observed in specific STs. For example, co-occurrences related to resistance phenotypes are found through the different STs, but are particularly evident in the MDR-associated ST167 and ST648 (52; 53). ST-specific differences that confer pathogenicity and resistance in *E. coli* are well-studied (43; 53; 54), but this work provides a new layer of understanding into how the accessory genome may interact to this end. We suggest that certain gene collections are required by specific lineages and that this may be driven in part by MGEs.

Alongside these gene co-occurrences of known function, we have also uncovered a network of genes with unknown function that influence the structure of the pangenome through the high number of genes that they co-occur with and avoid. *E. coli* has fewer genes of unknown function than most prokaryotes, although there is evidence that many of the accessory genes in lineages such as ST131 are of hypothetical function (40). The number of coincident gene relationships formed by such genes here highlights the challenges in understanding the global prokaryote pangenome. Given the prevalence of MGEs in the co-occurrence network, it is tempting to conclude that at least some of the high proportion of co-occurrence hub genes identified as encoding hypothetical proteins may be related to mobility.

The data presented here support the concept that pangenomes create pangenomes. The diversity of gene content in a cosmopolitan species such as *E. coli* means that the fitness effect of gaining and losing individual genes is not the same for all constituent genomes. We observed the presence or absence of some genes only when other genes are also present or absent. We consistently observed some pairs of genes co-occurring or avoiding repeatedly across the diversity of the group of genomes, and root excluders consistently avoiding certain cliques. We observed unknown ORFs that significantly co-occur in the same genome, even though their distribution in the pangenome is patchy. There is therefore an emerging logic to the *E. coli* pangenome that clearly identifies natural selection to have frequently dominated over genetic drift.

## Acknowledgements

RJH was supported by the BBSRC (BB/N018044/2), awarded to JOM. FJW was funded by a Marie Skłodowska-Curie Individual Fellowship (GA no. 793818). EAC was funded by the Wellcome AAMR DTP and CC by the Wellcome Midas DTP. We thank M.R. Domingo-Sananes and S. Thorpe for their insightful feedback on the work presented here.

## Supplementary

**Fig. S1.**
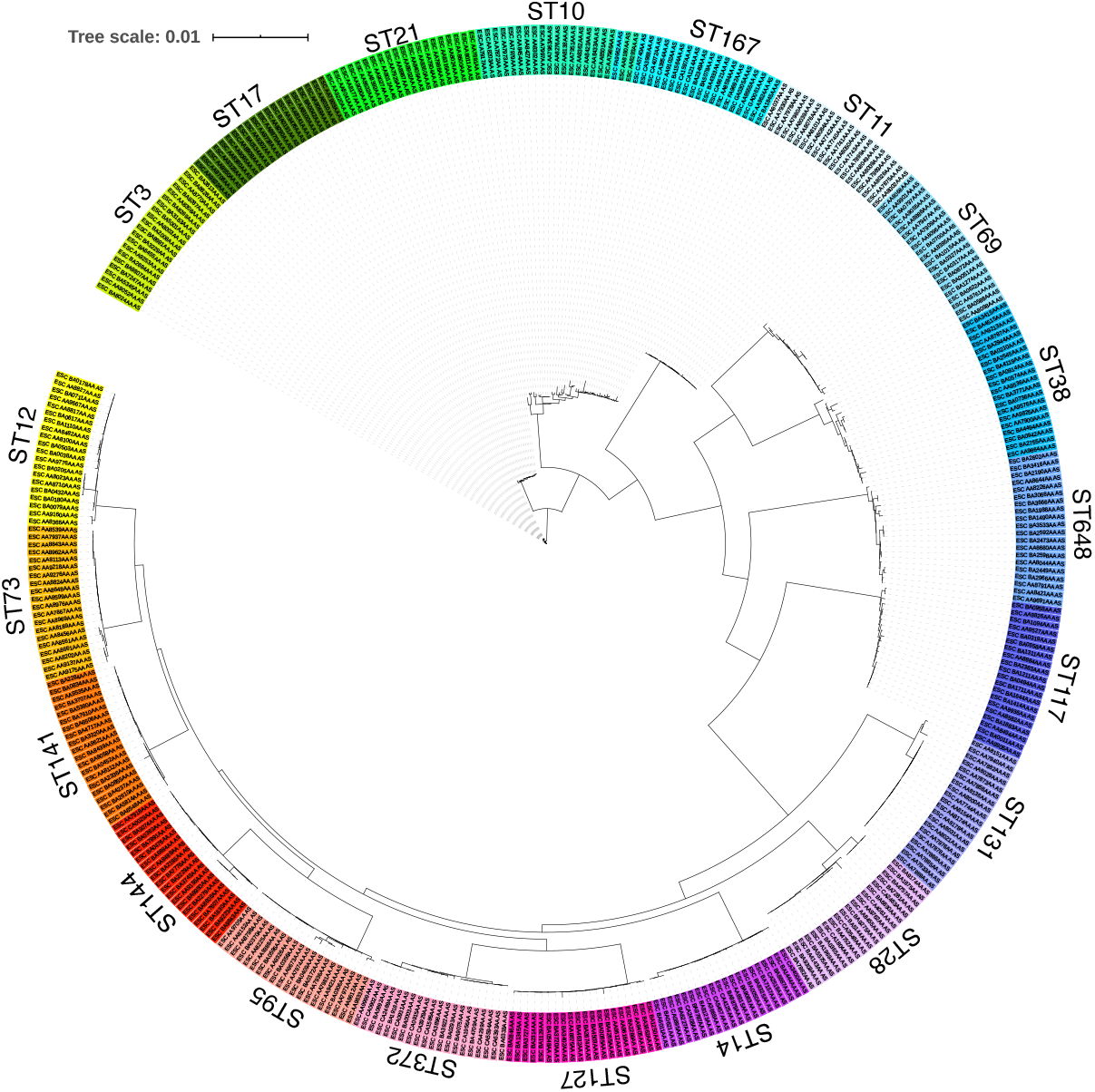
An *E. coli* core gene phylogeny. Phylogeny of the 400 genomes used in this study, constructed from the trimmed core gene alignment using IQ-Tree based on the GTR+I+G substitution model. Labels are coloured by ST.

**Fig. S2.**
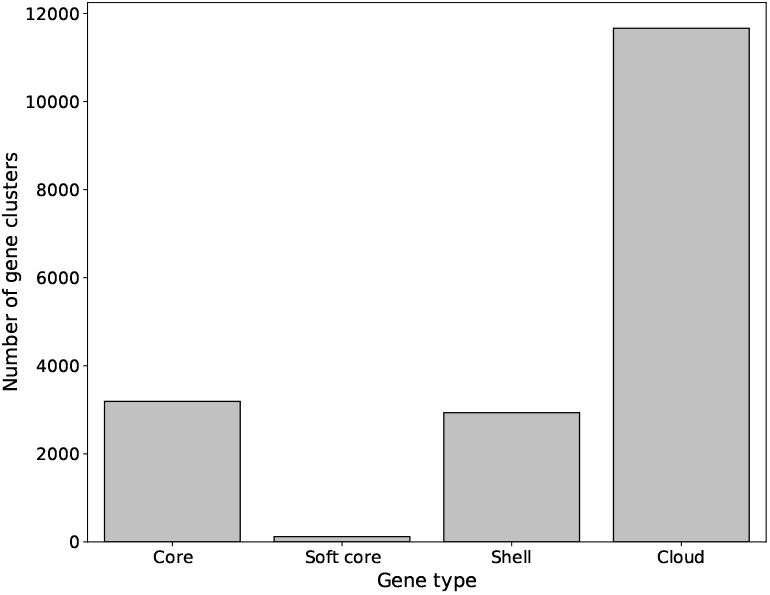
Distribution of core, soft core, shell, and cloud genes in this *E. coli* pangenome.

**Fig. S3.**
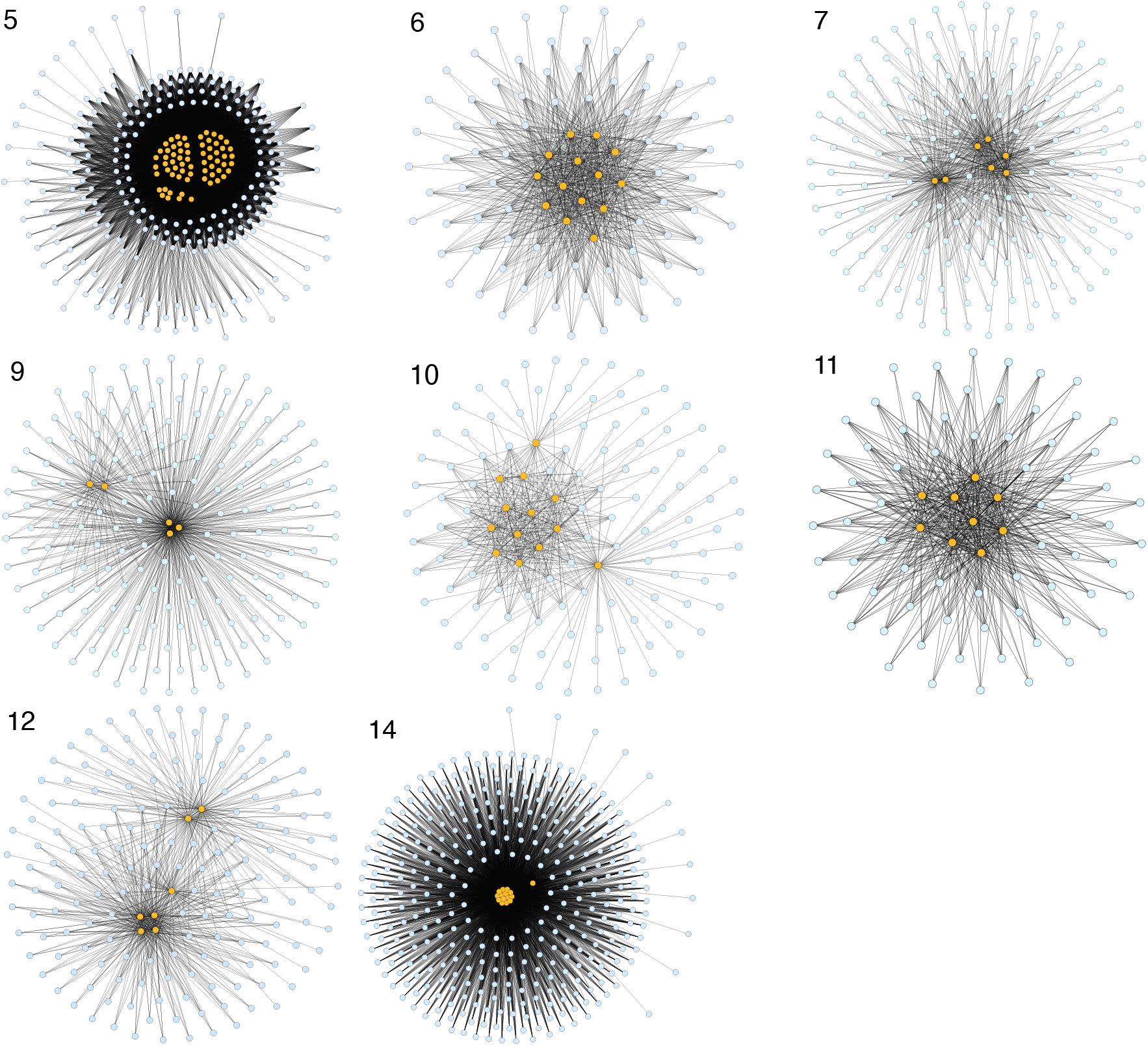
Gene relationships in avoidance connected components that contain a root excluder. Genes avoided by the root excluders (orange) co-occur with one another and with other genes in the accessory genome (blue). Networks numbered according to avoidance connected component in Fig. 1b and the Supplementary Data files.

**Table S1.**
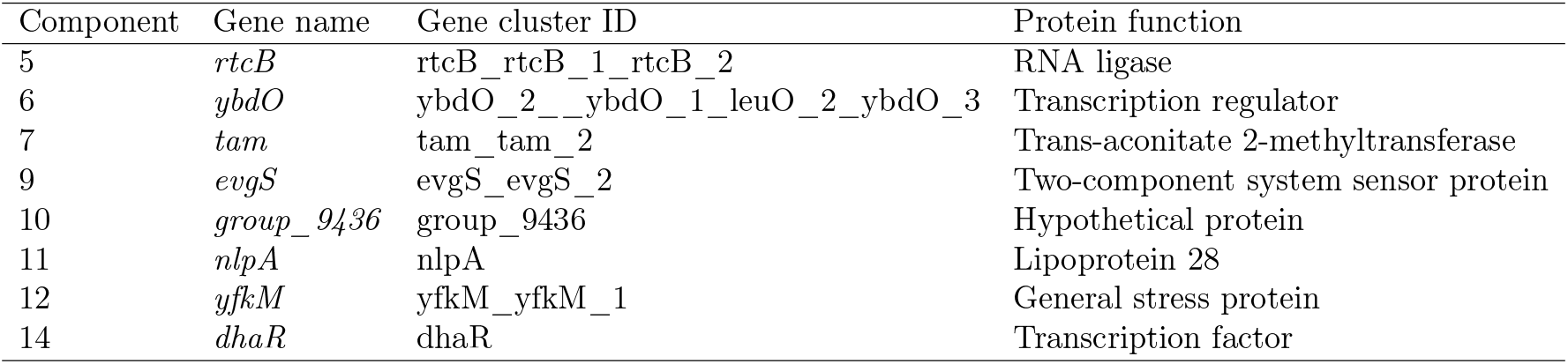
All root excluder genes and their corresponding avoidance connected component. Component number refers to that given in the Coinfinder output provided as Supplementary Data. The gene cluster IDs are given as in the Panaroo gene presence-absence matrix.

